# Method for isolation of small extracellular vesicles from different biofluids and workflow for Mass Spectrometry based-shotgun proteomics and RNA isolation

**DOI:** 10.1101/2024.05.15.594449

**Authors:** Pratibha Sharma, Rajinder K Dhamija

## Abstract

Small extracellular vesicles (sEVs) or exosomes are small-sized (30-150 nm), nanoparticles that are released from almost all cells under normal and pathophysiological conditions. The sEVs have a vital role in biological systems as they communicate and transfer their contents, such as proteins, lipids, and nucleic acids, from the cells of origin to nearby or distant cells. In recent years, there has been a growing interest in isolating sEVs for use in disease mechanisms, clinical diagnoses, and therapeutics. Due to their small size sEVs can be observed using electron microscopy. The size distribution and concentration were checked by Nanoparticle Tracking Analysis. Western blotting confirmed the presence of exosome markers. The ease of obtaining patient samples from biofluids like plasma, saliva, and urine makes them a valuable source for diagnostic purposes by isolating sEVs to diagnose and predict diseases early. However, there is no specific protocol to perform it altogether. We have developed an improved ultracentrifugation method using gradient ultracentrifugation and ultrafiltration, which results in higher sEVs purity and yield. We have tested this method on plasma, saliva, and urine at a single platform, and we have isolated proteins and RNA from exosomes for their downstream applications. Our method is simple to use and can be utilized for clinical research biomarker applications, in understanding disease mechanisms and monitoring its progressions from biofluid sample collections.

## Introduction

Biofluid sEVs or exosomes hold significant potential as specific disease cell information indicative of pathophysiological conditions [1-4]. However, the major challenge is the sample heterogeneity and common methodology to isolate and purify SEVs suitable for downstream applications [5, 6]. The sEVs are small-sized particles surrounded by lipid bilayers that are released by cells and play a vital role in cell-to-cell communication by transferring nucleotides, proteins, and lipids. They can be found in various bodily fluids and cell culture media and are being studied for their potential use as biomarkers and therapeutic agents in cancer and neurological disorders. Exosomes vary in size, content, function, and sources [2]. Early endosomes form multivesicular bodies (MVBs) that transport cargo to the extracellular space or for lysosomal degradation. The exosome formation and release are based on endosome sorting complexes required for the transport (ESCRT) pathway. Microvesicles (100-1000 nm) are extracellular vesicles formed by direct budding from plasma membrane proteins. Exosomes can be distinguished from microvesicles using exosome protein markers. Apoptotic bodies (800-5000 nm) are larger in size and similar in content to cell lysates. Despite variations in size, exosomes and microvesicles are studied for biomarker discovery, drug delivery, and therapeutics [7].

Various methods have been described, including ultracentrifugation, ultrafiltration, size-exclusion chromatography, Polyethylene glycol, immunocapture, and microfluidics [8]. Differential ultracentrifugation and density gradient ultracentrifugation methods are gold standards. A few additional steps from traditional methods will help provide purity and high yield. In our protocol, we have described the method for the isolation of sEVs from plasma, saliva, and urine from healthy human control. We have used electron microscopy-TEM and SEM to visualize sEVs. The particle size distribution and concentration are detected by Nanoparticle Tracking Analysis. The presence of exosome markers such as CD9, CD63, Flotilin 1, and TSG 101 has been checked by western blotting. We have further isolated the proteins and miRNA for its downstream applications. Our method can be performed with simple-to-follow steps and cost-effective manner from low to high volume of samples from various biofluids.

## Materials and Methods

Healthy human biofluids including blood plasma, saliva, and urine were used for our methods presented here. The overview of the improved method for the isolation of sEVs has been shown in Figure 1. This protocol can be used for other biofluids like cerebrospinal fluid (CSF), blood serum, milk, tears, semen, epididymal fluid, amniotic fluid, and synovial fluids as well as cell cultures from various tissues with some modifications at initial steps [9, 10].

**Figure 1.**
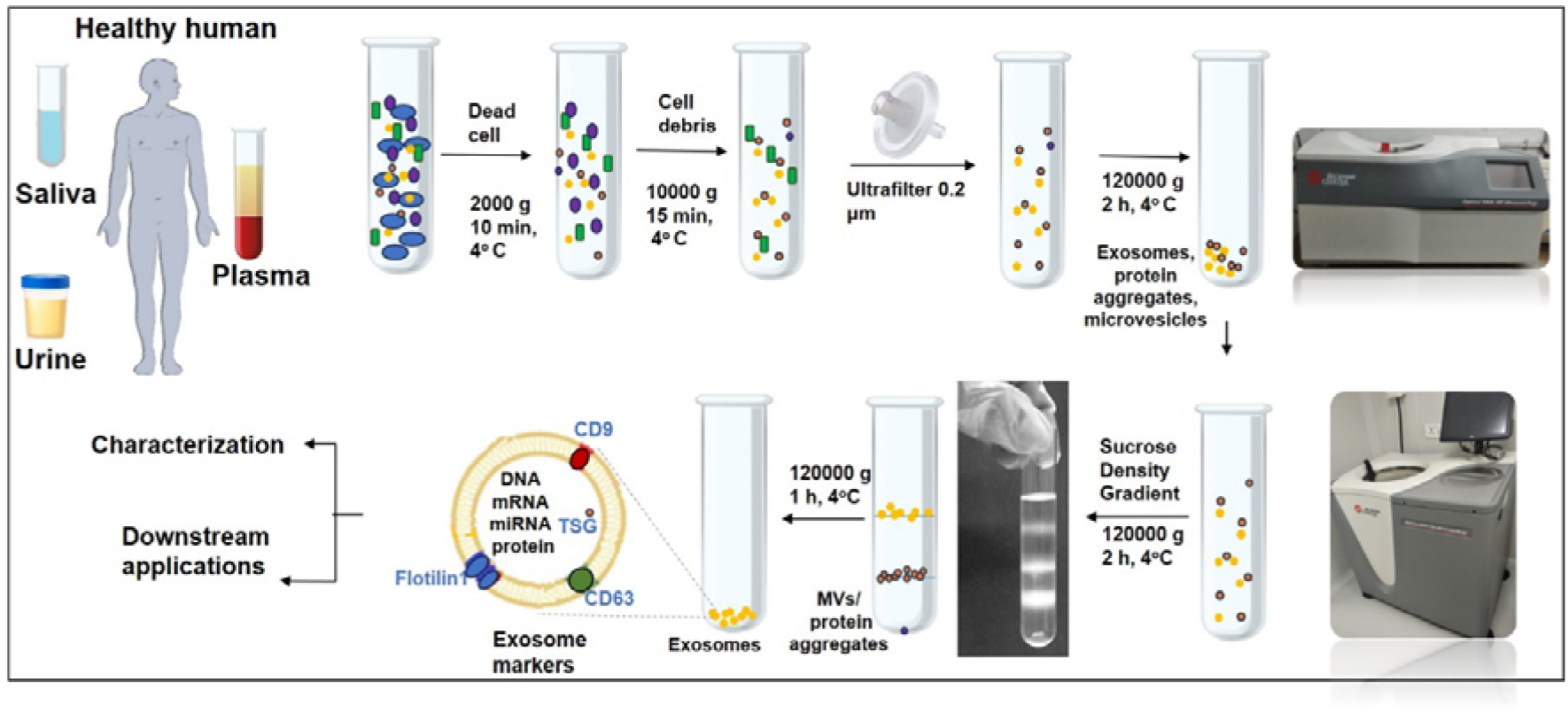
Overview of methodology for isolation of small extracellular vesicles or exosomes from plasma, saliva, and urine for downstream applications

### 1. Collection of samples

Blood (1 mL), saliva (2 mL), and urine (10 mL) were collected from healthy volunteers.

Note: The samples were collected in sterilized vacutainers and kept immediately on ice to prevent microbial contaminations and prevent proteolysis. Also, a protease inhibitor cocktail should be added for protein studies. This will help preserve the extracellular vesicle membrane proteins.

Note: Urine and saliva samples contain a larger amount of water (>90%) and salts in them. As a precautionary step, it is advisable to collect at least 2 mL of saliva and 10 mL of Urine for sEVs protein isolation for mass spectrometry and RNA isolations.

#### 1.1 Urine

1. Urine samples were collected into sterile urine vials (of volume 30 mL). The samples having a blood-like appearance were discarded. Note: We have collected the midstream urine sample (preferably 2^nd^ or 3^rd^ urinated samples). The biological appearance of a pale-yellow color, without blood contaminations, clear not cloudy, pH (4.5-8) was considered.
2. These were transferred to sterilized falcon tubes (15 mL, Corning, Inc.) and centrifuged at 2,000 g at 4°C for 15 min. Note: This step will remove the contaminants (bacteria, casts, and debris).
3. A clear supernatant leaving 1mL below obtained pelleted material was collected and stored in aliquots of 1.5 mL each in sterilized centrifuge tubes.

#### 1.2 Saliva

1. Unstipulated saliva samples were collected in sterilized tubes (50mL Falcon, Corning) immersed in ice. Note: Participants were refrained from eating (8 hours before collection) or drinking (30 minutes before collection). The biological characteristics of saliva samples were examined for color (transparent, no blood), turbidity (clear), and pH (>6.0) to ensure that there were no notable discrepancies. Volunteers with severe dental decays, oral bacterial infections, and oral malignancy were not used.
2. The samples were kept on ice undisturbed for 15 min. The upper clear material was pipetted into sterilized 1.5 mL Eppendorf tubes. Note: The submandibular gland produces saliva which is highly viscous and difficult to process technically.
3. The samples were centrifuged at 2,000 g for 10 minutes at 4°C to remove debris.
4. The samples were labeled and stored at -80°C.

#### 1.3 Plasma

1. Blood samples will be collected and transferred to the additive EDTA-treated (Lavender top) tubes.
2. These will be centrifuged at 1,500g for 10 min. at 4°C to remove cells.
3. Plasma aliquots in 1.5mL Eppendorf tubes will be stored at -80°C. Note: Care must be taken while collecting the plasma layer. Pipetting must not disturb the middle layer a whitish cloudy appearance and bottom layers containing mononuclear cells, other platelets, and red cells.

### 2. Methods for extracting small extracellular vesicles

1. Isolated plasma, saliva, and urine samples were centrifuged at 15,000 g for 15 min. at 4°C. Note: This step will remove the cell fragments and organelles. Note: For the -80°C frozen samples, these should be vortexed properly before melting completely. This step is necessary for the increase in exosome recovery.
2. Each sample supernatant will be filtered through Presterilized Polyethersulfone (PES) membrane syringe filters, size 0.2 µm (Merck Millipore). Note: Care must be taken during filtration of saliva samples. It may cause hindrance due to the viscous materials present. Although plasma and urine will be a little easier to filter. Note: The filtrate should be first filtered with distilled water. However, due to hold-up volumes, some samples may be lost during filtering.
3. Transfer each filtrate of plasma, saliva, and urine separately into ultra-clear tubes accordingly.
4. These will be diluted in sterile 0.1 M Phosphate buffered saline (PBS), pH-7.2 in sample to PBS ratio 1:3.
5. Ultracentrifuge at 1,20,000g for 2h at 4°C (Optima max instrument, Thermo Scientific).
6. The upper solution will be discarded leaving 100-150µL at the bottom of the tube. Note: Care must be taken to collect the bottom material. A very hazy small pellet will be seen sticking to the bottom of the tube. Carefully collect by pipetting up and down. Note: These will also include microvesicles and protein aggregates of similar sizes along with small extracellular vesicles. The additional steps are recommended for the isolation of small extracellular vesicles.

#### 2.2 Additional steps for purity in small extracellular vesicles

7 The collected samples from 6) will be repeated for dilution as mentioned in step 4).
8 Prepared sucrose density gradient solution from 1 M, 1.4 M, and 1.8 M in three layers. Overlay the obtained samples on top and centrifuge 1,20,000 g for 2 h at 4°C using swing rotors.
9 The topmost solution will be discarded. The small extracellular vesicles will be found at the interface between 1 M and 1.4 M. The rest bottom will be discarded as microvesicles. This will help remove protein aggregates. Note: Instead of three layers, the one-step method of preparing 1.4M sucrose in PBS can also be used. In this, the topmost layer will be collected above 1.4M sucrose, and the rest discarded. But this may include protein aggregates.
10 Collect exosome fraction and repeat ultracentrifugation by diluting as mentioned in step 4).
11 The samples will be collected in fresh 1.5 mL Eppendorf tubes, labeled, and stored at -80°C. These samples can be used immediately for downstream applications. Note: It is advisable to collect fresh plasma, saliva, and urine samples followed by immediate processing for the characterization by electron microscopy, flow cytometry, and particle size methods.

### 3. Sample preparation method for Electron microscopy

#### 3.1 Transmission electron microscopy

1. The samples collected will be fixed by resuspending in 2% paraformaldehyde, and 2.5% glutaraldehyde in 0.1 M phosphate buffer (PB) for 2 h at 4°C.
2. Washed twice by diluting in PB after each centrifuge at 10,000 g for 30 min at RT.
3. A 100 µL solution at the bottom was taken and diluted in 100 µL distilled water. The 50 µL of solution will be taken and incubated on a Formvar-coated copper grid (200 mesh) for 5 min.
4. Staining will be performed using 2% phosphotungstic acid, pH-7.0 for 15 sec., and blotted dry.
5. Samples can be viewed under the TEM instrument (Tecnai G2-20, Fei company, The Netherlands)

#### 3.2 Scanning electron microscopy

1. The freshly isolated plasma, saliva, and urine sEVs will be fixed in 4% paraformaldehyde and 2.5% glutaraldehyde in 0.1M PB, pH 7.2 at 4°C for one h.
2. Washed twice by diluting in distilled water after each centrifuge at 10,000 g for 30 min at RT.
3. Each 100 µL volume of sample was dissolved twice in distilled water.
4. Each 200 µL sample was mounted on the coverslip and allowed to dry naturally under a desiccator.
5. The dried coverslips were mounted over the metal stub and sputter coating with colloidal gold.
6. Samples can be viewed under SEM Zeiss Evo-18 microscope.

### 4. Particle Size experiment

1. The samples obtained after ultracentrifugation will be diluted in PBS as 500-4000 folds. An injection volume of 1 mL will be used. Note: Dilution in PBS is required to reduce the number of particles to less than 400/image. The instrument’s linear dynamic range is (∼ 10^7^- 10^9^ particles/mL; Zeta View NTA (Particle Matrix software version 8.05).
2. The samples will be viewed under 11 different positions at room temperature. Note: The samples can be used in the range from 1µL to 5µL obtained after the ultracentrifugation additional step. Plasma, saliva, and urine samples will have different concentrations of sEVs in them.

### 5. Protein isolation and preparation for western blotting

1. For isolation of protein Trichloroacetic acid (TCA)-acetone precipitation will be used. Note: TCA should not be used for targeted proteomics analysis of enzymes from exosomes. It might damage them. Note: The samples may be lost during this method. So, it is advisable to initially take an appropriate amount.
2. The pellet thus obtained will be dissolved in RIPA buffer (50mM Tris, pH 8.0, 150 mM NaCl, 1% NP-40, 0.5% Sodium deoxycholate, 0.1% SDS). Add PMSF as per sample volume. Vortex and spin briefly to avoid bubbles. Note: RIPA buffer needs additional steps in mass spectrometry for cleaning by Zip-tip or desalting. Note: The pellet obtained after the protein precipitation from plasma sEVs was difficult to dissolve, so a mild sonication with probe sonicator was used for 3 cycles each 10 sec ON/OFF for 2 min. Also, the samples may be dissolved in 5X Laemmli sample buffer.
3. The protein content of sEVs was determined using a BCA protein assay kit (Pierce, Thermo Scientific). Store at -80°C.
4. 20 µg of each sample will be used to run 12% SDS-PAGE gel. Western blotting for these samples can be performed.

### 6. Protein digestion, desalting and quantitation

1. For this initially 100 μg of each sample will be used. These will be diluted in 50 mM Ammonium Bicarbonate buffer. Note: For lysis buffer, the urea concentration must be less than 1M. Dilute with ammonium bicarbonate according to checking pH ≥ 8 using pH paper strips (Sigma-Aldrich).
2. 20 mM Tris (2-carboxyethyl) phosphine (TCEP, Sigma-Aldrich) will be used to reduce disulfide bonds in proteins. Incubate at 37°C for 1h.
3. Perform alkylation with 50 mM iodoacetamide (Sigma-Aldrich) for one hour at room temperature in the dark.
4. Add trypsin (V5280 Promega) to protein ratio was 1:20 and samples will be incubated at 37°C for 16 hours. The reaction was stopped using a final concentration of 1% Formic acid in the samples.
5. Dry the digested peptides using a vacuum concentrator. Note: The above method can be used for Label-free quantitation as well as iTRAQ or TMT label-based quantitation. Both methods need different setups in LC-MS/MS run and analysis.
6. Peptide desalting will be performed using peptide desalting spin columns (Pierce, Thermo Scientific) as per manufacturer protocol.
7. The dried peptides will be resuspended in 0.1% Formic acid.
8. Each 2 µL of sample peptides will be used to quantify peptide concentration with a Nanodrop instrument at absorbance 205 nm and 280 nm. The concentration in µg/ µL will be calculated.

### 7. LC-MS/MS and data analysis

1. For label-free quantitation, 2 µg of peptide was diluted in 10 µL of 0.1% FA.
2. For LC parameters, equilibrate the column and analytical column with 0.1% FA. The samples will be kept in an autosampler. 1 µg of digested peptide will be injected into the column.
3. The LC gradient program will be set according to the sample complexity using gradients for 125 min. for these samples.
4. MS parameters will be optimized using standard peptides. After optimization run the program.
5. The raw datafiles of the MS/MS spectrum obtained will be analyzed using commercially available MS protein analysis software.

### 8. Total RNA isolation

1. A kit by Invitrogen for Total RNA and protein isolation can be used. We have used this kit after the isolation of sEVs as discussed above. Note: For work with RNA clean-up of pipettes, a lab workbench with RNase decontaminating solution is necessary.
2. 100 µL of samples obtained from the additional step of exosome isolation for plasma, saliva, and urine samples will be diluted in 0.1M PBS to make the volume of 200 µL.
3. Add 100 µL of 2X Denaturing solution on ice and incubate for 5 min.
4. Add 100 µL of Acid-phenol: Chloroform to each sample. Mix thoroughly by vortex and centrifuge 10,000g for 5 min at RT. Note: Layers of the aqueous phase, interphase, and bottom organic phase will be seen clearly.
5. Carefully pipette the aqueous layer at the top in an RNase-free fresh tube.
6. Add 1.25 µL of 100% ethanol to 100 µL of each volume recovered at step 5.
7. Add mixture to filter cartridge and centrifuge at 10,000g for 20s. Allow the mixture to pass through completely and discard the filtrate.
8. Wash thrice using 500 µL of PBS for each sample.
9. Elute the RNA using 25 µL of nuclease-free water and repeat this step.
10. Collect and store the 50 µL volume of total RNA isolated from each sample, label, and store at -20°C.
11. The RNA concentration can be determined using a Nanodrop instrument. Note: The RNA yield will vary for different biofluids. The experiments may be repeated to get the requisite concentration. The collected total RNA sample at step 11, can be used further for isolation of small RNAs.

### Results and Discussions

We have presented a simple and reproducible methodology for isolating sEVs or exosomes from plasma, saliva, and urine of healthy human control. However, the number, concentration, and morphology of sEVs may vary depending on various factors such as the organism, age, gender, cell of origin, and disease condition [11]. Additionally, the physical state and storage conditions, such as temperature, buffer, pH, and duration, may also affect sEVs [12, 13].

#### Electron microscopy

We conducted TEM, SEM, and NTA analyses using fresh isolates [Figure 2]. We used 1 μl of plasma sEVs and 2 μl each of urine and saliva obtained after isolation and negative staining. During TEM analysis, we observed different sizes of sEVs ranging from 30-150 nm and overall round-shaped morphology. Plasma sEVs had more mixed sizes and numbers than saliva and urine sEVs. Salivary sEVs were observed to be close to each other, while urine sEVs were scattered and separated. Overall, the three samples from the healthy control were dark, round-like shaped, and uniform in size. However, the heterogeneity may vary during diseased conditions as shown [14]. SEM analysis observed uniformly scattered, round-shaped, shiny, light-colored plasma and urine sEVs. Saliva sEVs were round and closer to each other rather than scattered. These represented blebbing with an internal membrane. The range of sizes varied largely in plasma sEVs mean of 76±52 nm. The sEVs <100nm sizes were largely observed. The saliva sEVs were mostly from size 100-150nm, mean of 120±7 nm. The urine sEVs were larger in their surface size from 150-200 nm, a mean of 150±8 nm. A small proportion of 3% was observed having larger diameters up to 215 nm in urine sEVs. These surface topologies observed can be useful for understanding biological processes through sEVs interactions.

**Figure 2.**
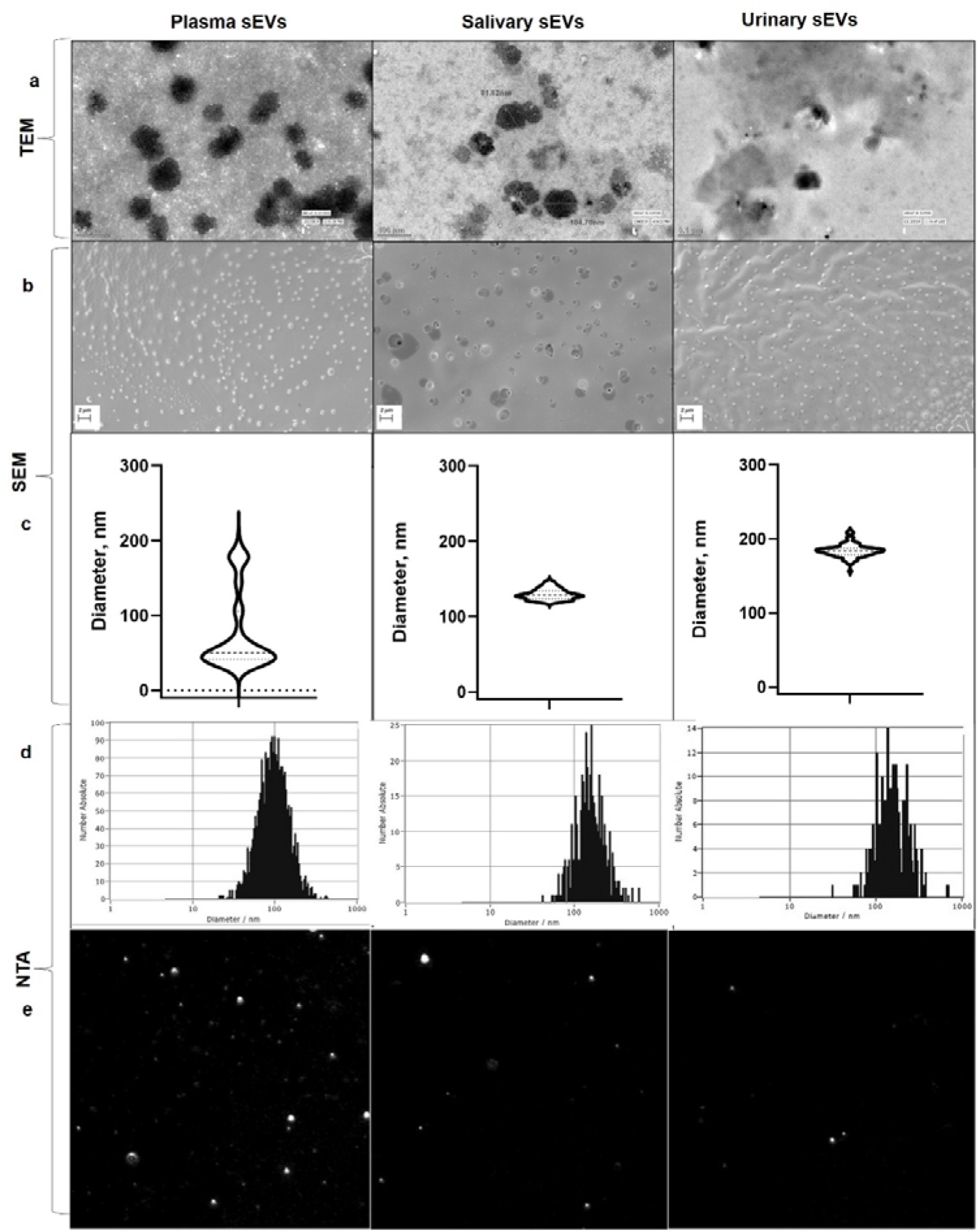
Characterization of sEVs from plasma, saliva, and urine. a) Transmission electron microscopy analysis of each sample showing representative images. b) Scanning electron microscopy surface morphology of each sample showing representative images. c) SEM obtained size of sEVs surfaces as analyzed using image J for each sample (diameter in nm). d) Nanoparticle tracking analysis histogram for plasma, saliva, and urine sEVs concentration and size distributions. e) Representative screenshots of particles as detected using NTA software for each sample.

#### Nano-tracking particle analysis

We conducted a Nano-tracking particle analysis to determine the size distribution and concentration of sEVs in three different samples. We used 1 µL plasma, 2 µL saliva, and 2 µL urine sEVs diluted in 1 mL PBS for the analysis. The results showed that the concentration of sEVs was higher in plasma compared to saliva and urine samples. Furthermore, we observed more diverse populations in plasma with a greater number of small-sized sEVs mean 140±78 nm, 2.7x10^11^ Particles/ mL. On the other hand, the saliva and urine sEVs were larger in diameter and lower in concentration, with sizes of 174±70 nm 1.4x10^10^ Particles/ mL and 170±79 nm, 1.5x10^10^ Particles/ mL respectively. We also noticed that the sizes in diameters that appeared in NTA were larger compared to TEM. This may be due to their dynamics in liquid and the Brownian motion of particles that they appeared bigger than original size. NTA provides specific data in the form of size distribution and concentration, while the accurate size can be determined from TEM. Further, we performed western blotting using exosome markers such as CD63, CD9, Flotilin-1, and TSG 101, which have been described and published previously [4, 14]. For this, we isolated proteins after the TCA-Acetone protein precipitation method.

#### Protein isolation, SDS PAGE, and western blotting

After characterizing the sEVs, we performed SDS-PAGE on the protein samples [Figure 3a-b]. The plasma pellet obtained after isolating sEVs may be sticky, so you must be careful during trituration and avoid sample loss. Compared to the saliva and urine sEVs, the plasma sEVs had a larger pellet of light pale color that may be difficult to dissolve. To help with this, we suggest using RIPA buffer or mild sonication with a probe sonicator in PBS buffer or incubating at 37°C for 5 minutes. The saliva sEVs pellet may be viscous and have an opaque color. However, the urine sEVs were non-sticky and easy to dissolve in any buffer. You may use a lysis buffer as well. For SDS PAGE, we have used 20 μg each of plasma, saliva, and urine sEVs protein samples to run 12% of SDS-PAGE gel. We observed that urine sEVs had a greater number of high molecular weight proteins [4]. A much diverse range of saliva sEV proteins were observed from both high and low molecular weights. Further, the proteins were checked by western blotting for the presence of exosome protein markers TSG 101 and Flotilin 1 [Figure 4]. We have previously presented other gel images using CD 63, and CD9 exosome markers [4, 14].

**Figure 3.**
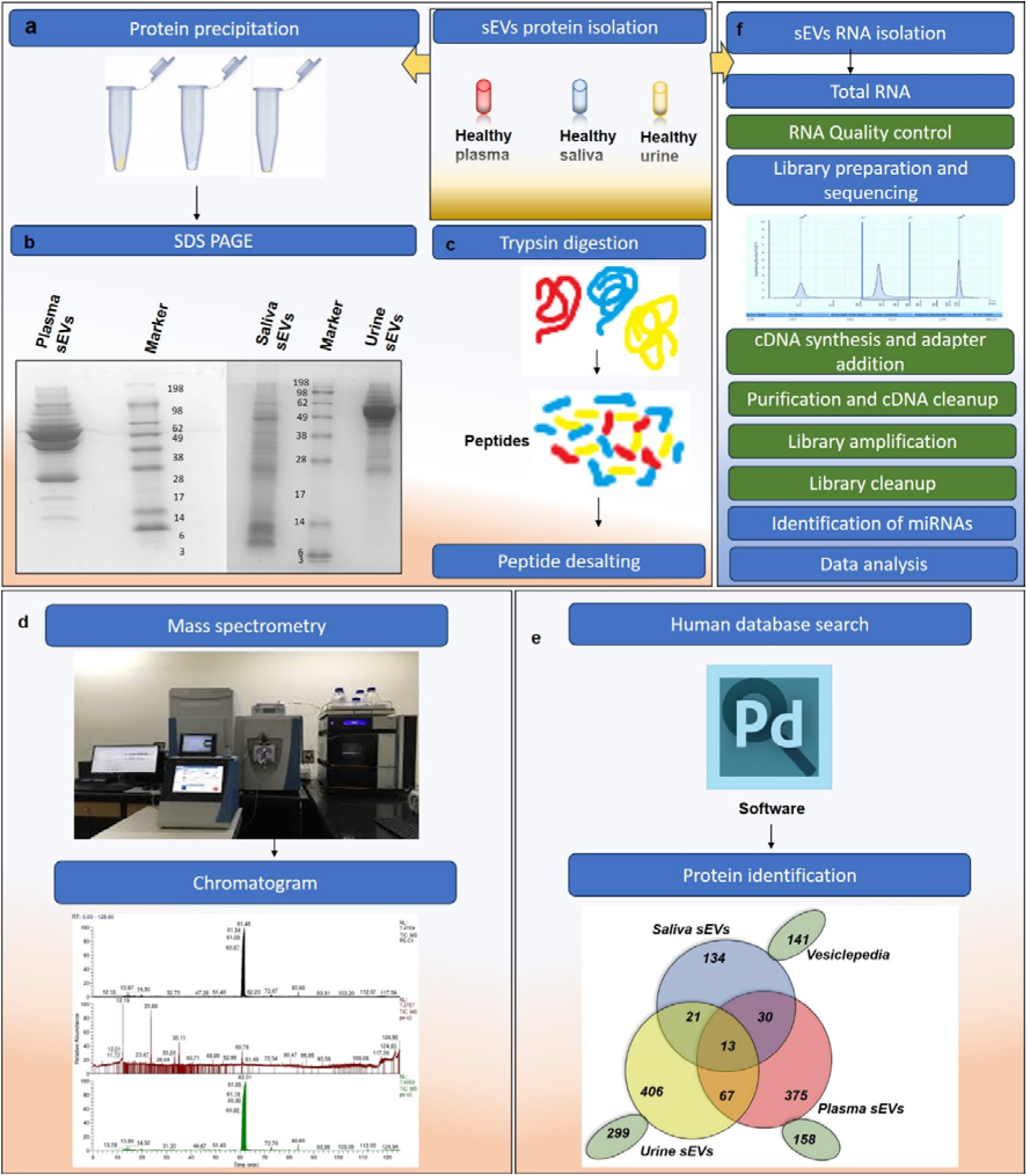
The isolation of protein and RNA from plasma, saliva, and urine for downstream applications. a) protein precipitation from each sample sEVs. b) 12% SDS PAGE of plasma, saliva, and urine sEVs proteins of 20 μg each, 8 μL pre-stained marker, gel stained with colloidal Coomassie stain. c) In-solution trypsin digestion of 50 μg plasma sEVs protein and 100 μg each of saliva and urine sEVs protein, desalted and concentrated by speed vac. d) Chromatogram obtained after mass spectrometry analysis by Orbitrap instrument. e) Proteome discoverer software analysis for the number of proteins identified from the Uniprot database searched against specie *Homo sapiens*. f) Isolation of total RNA and miRNA for further applications.

**Figure 4:**
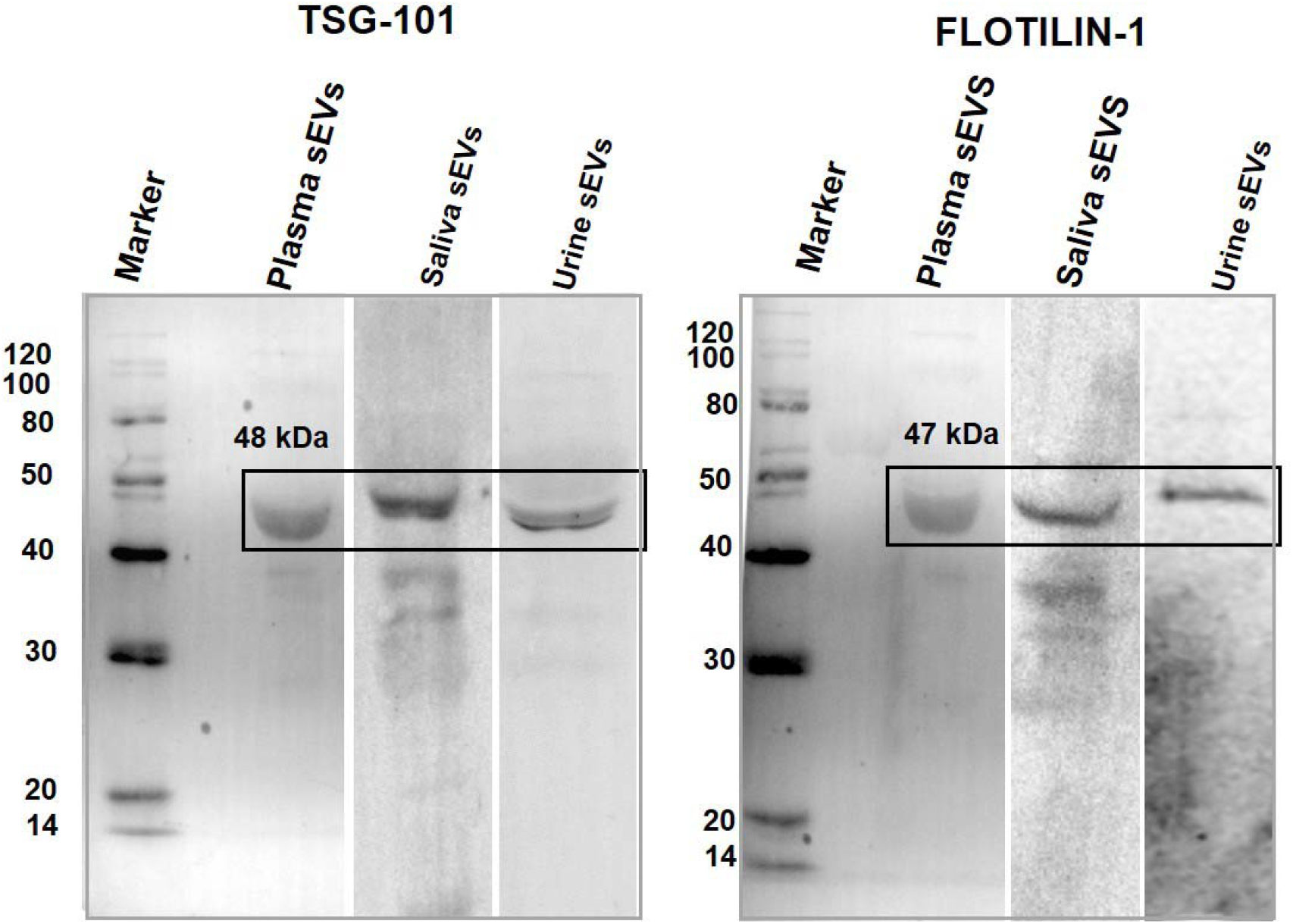
Characterization of plasma, saliva, and urine sEVs using western blotting markers. Western blotting for exosome markers Anti-TSG-101 and Anti-Flotilin-1 in healthy control plasma sEVs, saliva sEVs and urine sEVs each lane of 20 μg, Secondary Ab-Goat anti-rabbit IgG H&L; developed using ECL. Marker- a mixture of Magic Mark XP WB & See blue plus2 pre-stained protein standard; Image using ECL Azure biosystems.

#### Downstream applications using protein and RNA

Protein and RNA analysis is essential for studying various diseases. To perform the analysis, we used LC-MS/MS mass spectrometry with Orbitrap and searched the chromatograms using Proteome Discoverer software. We identified a qualitative number of proteins from plasma sEVs (n=485), saliva sEVs (n=198), and urine sEVs (n=507), as depicted in Figure 3c-e. Further, we cross-checked the identified proteins with the Vesiclepedia [15], which is a manually curated data containing proteins from various extracellular vesicles and exosomes. We found that approximately 62 percent of the identified proteins were previously reported from sEVs in these biofluids. The identified proteins in comparison to diseased conditions and their differential expressions may be useful to study as clinical biomarkers from biofluids. The biomarker discovery from sEVs can be done using total RNA as well as small non-coding RNA, as shown in the workflow depicted in Figure 3f.

The proteins from plasma had lower expression of albumin protein from sEVs [Figure 3b], which is generally found abundant in blood samples. This makes it suitable for mass spectrometry-based proteomic biomarker research, where we need to deplete such proteins to unmask the low molecular weight proteins. However, when using sEVs, depletion of such proteins may not be required while using sEVs. These sEVs lack proteolysis and are stable due to being bound by the plasma membrane. The method we have described does not require the pre-depletion of abundant plasma proteins, contrary to another author’s publication [16]. We have identified 485 proteins from plasma without such depletion, identified by mass spectrometry. Our method with combined approaches will be useful in sEVs isolation and further applications.

Exosomes are implicated in spreading diseases like cancer, metabolic disorders, atherosclerosis, cardiovascular diseases, and neurological disorders. The purpose of presenting this protocol is to provide further support to the clinical biomarker field. By using a multi-omics approach, which involves identifying biomarkers from sources such as DNA, RNA, protein, and lipids, we can obtain more comprehensive and beneficial results from non-invasive sources such as blood, urine, and saliva. Our cost-effective and easy-to-follow methodology will be helpful in this regard. Our approach can be completed in less than 6 hours, depending on the instrument used. Ultracentrifugation is a widely accepted protocol for isolating sEVs or exosomes. Our method, which includes additional steps of gradient ultracentrifugation, can work with a range of low to high-viscosity samples, such as saliva, to provide enhanced purity in the isolation of sEVs. Along with the ultrafiltration step, which can be used for large sample volumes, we can achieve a higher purity and yield. The same protocol can be used to isolate sEVs from both healthy and diseased cell lines using exosome-free FBS media [17] Our modified method for biofluids may have extensive applications in the field of clinical biomarker discovery and research depending on the research type.

## Funding

Our study was supported by fellowship grant number R.12013/07/2019-HR from the Department of Health Research, Ministry of Health & Family Welfare (MoHFW), India.

## Acknowledgments

We would like to acknowledge SAIF, AIIMS for Electron Microscopy and NTA, Department of Biochemistry for Ultracentrifugation and other basic lab facility, and SAIF, IIT Bombay for mass spectrometry work. Our study involves the use of healthy control samples which is supported by ethical committee approval with reference number IEC/425/07.06.2019, RP-21/2019 from the Institutional Ethics Committee, All India Institute of Medical Sciences, New Delhi, India. We would like to acknowledge the IHBAS facility for completing our manuscript.

## Notes

### Competing Interest Statement

The authors have declared no competing interest.

