## Supplementary material for "Method for isolation of small extracellular vesicles from different biofluids and workflow for Mass Spectrometry based-shotgun proteomics and RNA isolation": Supplementary information data - Copy.docx

**SUPPLEMENTARY DATA**

**Protein estimation and SDS PAGE of sEVs proteins isolated from plasma, saliva ad urine of healthy control by BCA Method**

|  |  |
| --- | --- |
| 25 | 0.018 |
| 125 | 0.10145 |
| 250 | 0.1937 |
| 500 | 0.33335 |
| 750 | 0.5186 |
| 1000 | 0.6495 |
| 1500 | 0.899 |
| 2000 | 1.1507 |

Each pellet obtained after density gradient ultracentrifugation as shown in Figure 1 was dissolved in 100 ul PBS and protein quantified by BCA protein estimation method. We obtained a concentration of plasma sEVs - approximately 6 μg/μL, saliva sEVs - μg/μL, and urine sEVs - μg/μL. The protein quantification results were also checked from the Nanodrop instrument and obtained the same protein concentrations.

-----

Western blotting

Primary

- antibody anti-human TSG101 (ab30871, Abcam, Cambridge, UK), dilution 1:1000
- anti-human flotillin-1 (ab41927, Abcam, Cambridge, UK), dilution 1:1000

Secondary

- anti-Rabbit IgG H&L (ab205718, Abcam Cambridge, UK), dilution 1: 5,000

Developed by exposure to Enhanced chemiluminescence (ECL) plus Western blotting detection reagents (Amersham Biosciences, USA). Images were obtained from the Azure systems 400 (Azure Biosystems, California, USA).
